# Prevalence of Antibiotic Resistance in Commensal Escherichia Coli among the Children in Rural Hill Communities of North East India

**DOI:** 10.1101/280198

**Authors:** Ashish Kumar Singh, Saurav Das, Samer Singh, Varsha Rani Gajamer, Nilu Pradhan, Yangchen Doma Lepcha, Hare Krishna Tiwari

## Abstract

Commensal bacteria are the representative of the reservoir of antibiotic resistance genes present in a community. Merely a few community-based studies on the prevalence of antibiotic resistance in commensal bacteria have been conducted so far in Southeast Asia and other parts of India. Northeastern India is still untapped regarding the surveillance of antibiotic-resistant genes and prevalence in commensal bacteria. In the present work, the prevalence of antibiotic resistance in commensal *Escherichia coli* was investigated along with the associated demographic factors in pre-school and school going children in rural areas of Sikkim. A total of 550 fecal *E. coli* isolates were obtained from children of the age 1-14 years living in different villages at various altitudes of Sikkim from July 2015 to June 2017. Standard antibiotic susceptibility testing of these isolates was performed. A structured questionnaire was designed to study the factors associated with carriage of antibiotic resistance in commensal *E. coli* isolates among children. Descriptive statistics analysis and a logistic regression model were used to identify the effect of external factors on antibiotic resistance pattern. High prevalence of resistance was found against commonly used antibiotics ampicillin (92%), ceftazidime (90%), cefoxitin (88%), streptomycin (40%) and tetracycline (36%) among the samples examined in our present study. No resistance to chloramphenicol was recorded. Fifty-two percent of the isolates were resistant to the combination of penicillin and quinolone group of antibiotics. Children living in nuclear families showed higher incidence of resistance to ampicillin (63.15%, OR 0.18,95% CI:0.11 – 0.28, p<0.01) while children of mothers having education up to school level displayed higher incidence of ceftazidime (59.27% OR 0.75, 95% CI:0.55 - 1.02, p<0.02). Our study demonstrates a high prevalence of antibiotic-resistant commensal *E. coli* against the commonly used antibiotics among children in the study area. A close association between different demographic factors and the pattern of carriage of antibiotic-resistant isolates was observed suggesting a concern over misuse of antibiotics and warrants a future threat of emerging multidrug resistant isolates.

## Introduction

During the last decade, an alarming worldwide increase in the incidence of community-acquired infections with bacteria resistant to multiple antibiotics of common use has been observed [1]. Use of antibiotics plays a crucial role in the development of antibiotic resistance [2]. All over the world, development of resistance to antibiotics is on the rise amongst pathogenic bacteria [3]. However, to counter the situation, very few new antibiotics have come into use in the last three decades [4]. Rising resistance to antimicrobials in pathogens is a worldwide problem and is particularly serious in developing countries, where alternative antimicrobials are often not available or too expensive [5]. Inappropriate use of antimicrobials is considered to be one of the main factors responsible for the high prevalence of antibiotic resistance in developing countries [5]. Increased antibiotic resistance in pathogens leads to increased mortality and morbidity, enhanced transmission of antibiotic-resistant bacteria and increased health costs [6].

A large number of commensal bacteria colonize the gastrointestinal tract of mammals [7]. The commensal bacteria reside in the gut without being eliminated, they play an important role in human nutrition and health, by promoting nutrient supply, preventing pathogen colonization, shaping and more importantly maintaining the homeostasis of the intestinal immune system [6]. Use of antibiotics affects the population of commensal bacteria in gut [5][8]. In community settings, young children tend to be the most exposed to antibiotics and several studies have revealed that younger children have the highest risk of carrying antibiotic resistant commensal bacteria [9][8][10].

*Escherichia coli* is a member of the *Enterobacteriaceae* family, the enteric bacteria, which are facultative anaerobic Gram-negative bacteria, commonly found in the intestinal tract of warm-blooded animals including humans [7]. *Escherichia coli*, a near-ubiquitous colonizer of the gastrointestinal tract in children and adults has often been used in studies of the incidence of antibiotic resistance in commensal bacteria [11]. In recent years, the potential role of the commensal microbiota for the emergence and spread of antimicrobial resistance in pathogens has been universally acknowledged [12]. Some species of the commensal microbiota, such as fecal *E. coli*, have been exploited as sensitive indicators in surveillance and spread of antimicrobial resistance [13].

India is among the nations with the highest burden of bacterial infections and the crude mortality from the infectious diseases (In India 417 per 100,000 persons dies due to bacterial infections) [14]. In 2010, India was the world’s largest consumer of antibiotics for human health with 12.9 x 10^9^ units of antibiotic consumption (∼10.7 units per person) [2]. Global antibiotic consumption index from 2000 - 2010 was high among the BRICS countries i.e., Brazil, Russia, India, China, and South Africa. Of the BRICS countries, 23% of the retail antibiotic sales was attributable to India [15]. Antimicrobial resistance in pathogens is a major public health concern in India. The emergence of antimicrobial resistance is not only limited to the older and more frequently used classes of drugs but there has also been a rapid increase in the emergence of resistance to the newer and more expensive drugs, like carbapenem [14]. Extended-spectrum beta-lactamase (ESBL) producing strains of *Enterobacteriaceae* have emerged as a big challenge in the hospitalized patients as well as in the community and studies have indicated that 61% of *E. coli* isolates are ESBL producers [14].

Limited research has been done with regards to prevalence of antibiotic resistance in *E. coli* isolates among Indian children [8][16][17]. However, none of these studies were done in the north eastern population, hence there is no data suggesting the prevalence of antibiotic resistance in this area. In the few studies conducted, the wide variation has been demonstrated in resistance pattern of *E. coli* isolates from reportedly healthy children [8]. To evaluate the correlation between increasing antibiotic use and the emergence of antibiotic-resistant pathogens, community-based surveillance could be of great use. The aim of the present study was to investigate the prevalence of antibiotic resistance in commensal *E. coli* harbored by children in the hills of northeastern Himalayan regions and to identify various demographic factors associated with its carriage or spread in the children.

## Materials and Methods

### Site and study duration

The selected study area was Sikkim state of India and the study was conducted during July 2015 to June 2017. Sikkim is a northeastern state of India surrounded by Tibetan Plateaus in North, Chumbi Valley of Tibet as well as Bhutan kingdom in East. South is surrounded by Darjeeling district of West Bengal and the West by the Kingdom of Nepal [18]. Sikkim has total population 6, 10,577 in which 5, 56,999 (91.22%) belongs to rural areas [19]. In Sikkim, the literacy rate is 81.4%. As compare to females (75.6), males (86.6) are more literate [20]. The land of Sikkim is divided into lower hills (Altitude 270 to 1500 meters), middle hills (Altitude 1500 to 2000 meters) and higher hills (Altitude 2000 to 3000 meters) based on the geographical parameter. Sikkim has an Alpine zone (Altitude above 3900 meters with vegetation) and Snowbound Land (Very high mountains without vegetation up to 8580 meters) [18].

### Survey

Sikkim has a total of 451 villages with a population of 4,55,962 as per census 2011[20]. The survey was carried out in randomly selected 150 villages of Sikkim at different altitude with a total average population of children was 4500. The study design and the demographic details of respondent were summarized in **Fig 1** and **Fig 2**. The Gram Panchayat Units (GPU) were communicated and given prior information about the survey to be conducted. The healthy children from age 1-14 years were included in this study (Healthy: children who are metabolically active without any symptomatic diseases especially gastrointestinal abnormalities like diarrhea, vomiting, abdominal cramps, nausea at the time of survey; Unhealthy: Children with gastrointestinal abnormalities at the time of survey). Unhealthy children were excluded. The minimum required sample size (n= 350) was calculated using the OpenEpi software version 3.0.1 with finite population correlation (FPC) factor having 95% confidence interval with frequency of resistant to at least one drug was 50% [21]. Prior to the study, proper consent was taken from the parents/guardians (**Supplementary File. S1**). The structured questionnaires were used to interview the parents for making the decision about the inclusion of their wards in the study **(Supplementary File. S2)**. They were invited to participate in the study with an assurance, if they found any difficulty regarding the survey, they can withdraw at any stage of the study and their details.

**Fig 1:**
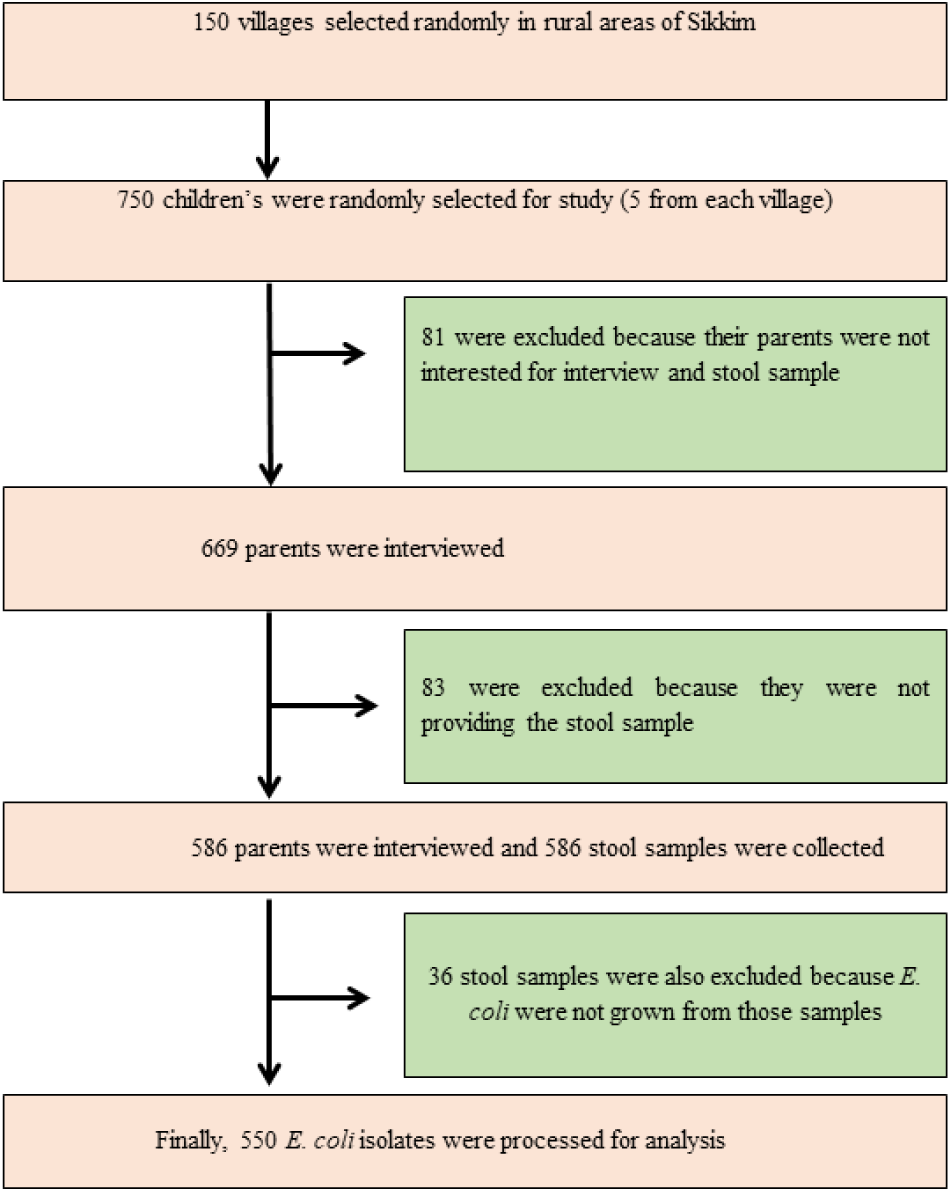
Study Design. Consents from parents/guardians was procured. Total 586 stool samples were collected from children (1-14) of different age and genders. Of the 586 stool samples, 36 stool samples were excluded as no *E. coli* isolates could be cultured from those samples. A total of 550 *E. coli* was isolated from 550 samples and used for the analysis.

**Fig 2:**
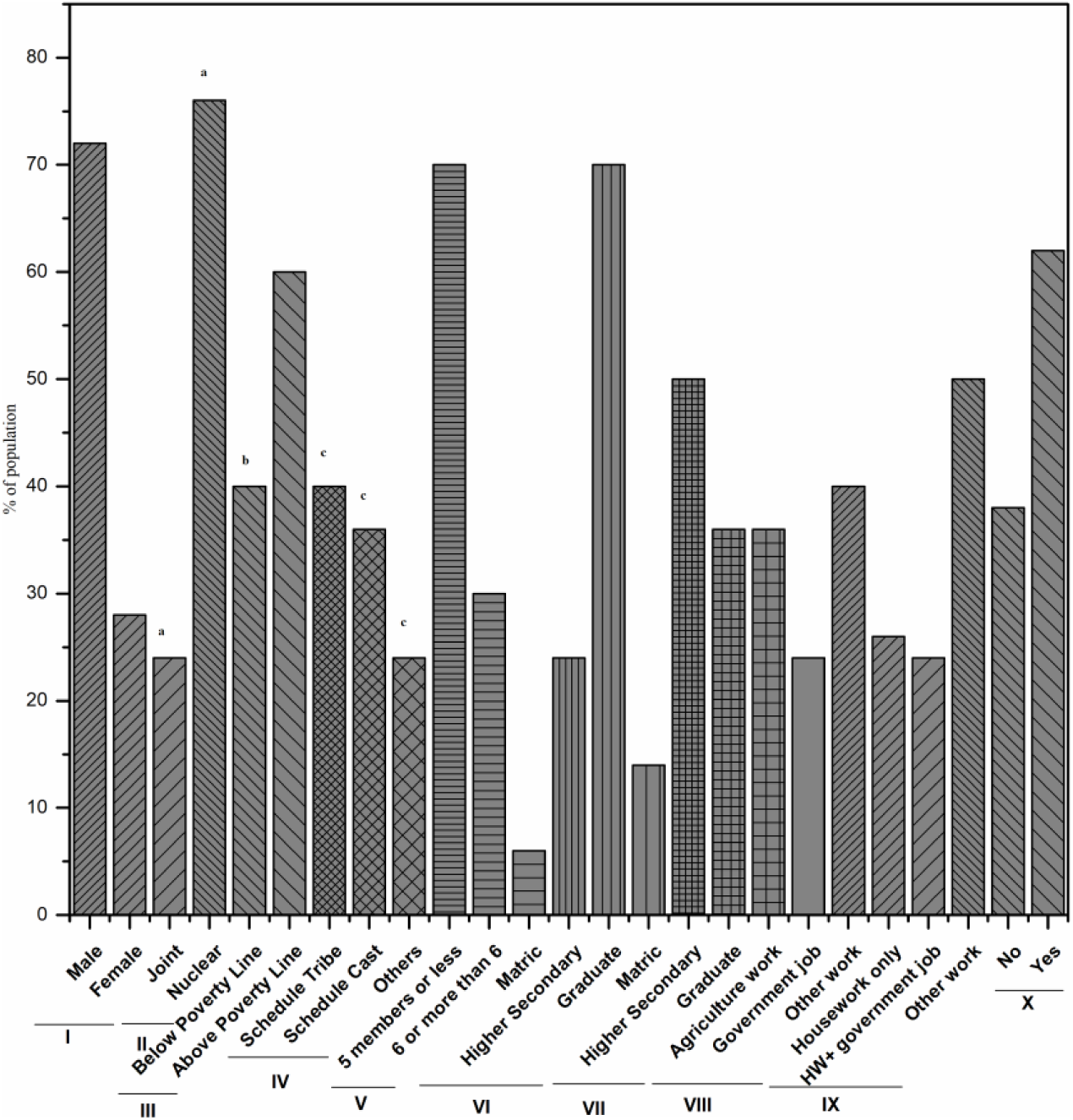
Demographic details of Community Participants. Demographic details of community participants and their families. Demographic details of community participants and their families are represented as a percentage of the population against the total population (N = 550). a = Nuclear family were referred to those family having single parents and children living on one household premises and Joint family referred to the family having parents, children and their relatives living in one household; b = Family below poverty line are referred to those family having possession of “Below Poverty Line (BPL)’ card issued by Government of India (GOI); c = Scheduled castes, backward castes and scheduled tribes are socially and economically deprived classes who were different social disadvantages from history. This special status of the group was issued by the GOI. The roman numeric stand for I= Sex of Children, II= Family type, III= Economic status, IV= Caste, V= Number of family members, VI= Paternal education, VII= Maternal education, VIII= Maternal occupation, IX= Antibiotics used last month.

### Fecal Sample Collection

During the survey, families were provided with clean, sterile, wide mouth, screw-capped specimen container for stool collection. Parents/guardians were instructed to collect morning stool [19]. All the stool samples were transported to the laboratory in cool condition on ice and processed within 6 hours. Samples were streaked on MacConkey agar (Hi-Media) and Eosin Methylene Blue (EMB) agar (Hi-media) for isolation of *E. coli*. *E. coli* isolates were confirmed by phenotypic study and standard biochemical tests [19].

The confirmed *E. coli* were further subjected to antibiotic susceptibility test by standard Kirby-Bauer disc diffusion method using CLSI guideline (Clinical and Laboratory Standards Institute, 2011) [22]. The antibiotics used in this study were ampicillin (30mcg), amoxicillin (30mcg), ciprofloxacin (5mcg), norfloxacin (10mcg), cefoxitin (30mcg), ceftazidime (30mcg), imipenem (10mcg), chloramphenicol (30mcg), streptomycin (10mcg), tetracycline (30mcg), netillin (30mcg), polymyxin-B (300U), ofloxacin (5mcg) and amikacin (30mcg).

A standard culture of *E. coli* (MTCC 10898) was used as a control with each batch of antimicrobial susceptibility test. The isolates showed resistant to three different groups of antibiotics were designated as multi-drug resistant bacteria. Extended-Spectrum ß-Lactamases test was also performed using ESBL Kit (Hi-Media, Mumbai, India). The *Klebsiella pneumoniae* (MTCC 9024) was used as positive control for ESBL detection.

### Statistical Analysis

Drug susceptibility data and filled questionnaires were collected and entered in XL-STAT. Questionnaires were analyzed as described previously [8]. Briefly, data were analyzed using descriptive statistics, bivariate analysis (cross-tabulation) and frequency. The significant level of p = 0.05 was used. The association between socio-demographic data and health behavior was correlated with the laboratory outcomes of *E. coli* resistance. The Association between the variables was tested using Chi-Square test. Those variables which approached statistical significance (p < 0.2) were entered into the multivariate logistic regression model. The odds ratio, confidence interval, and correlation between the antibiotic resistance pattern and the demographic structure (cluster analysis) were analyzed using R statistics (ver. 3.4, package = ggplot and epiR).

## Results

The demographic details of the families of 550 children from whom *E. coli* was isolated are mentioned in **Fig 2**. The median age of children which included in the study was 8.5 for male and 8 for female. Twenty-six percent (n = 143) of the children had a history of gastrointestinal illness in the last three weeks prior to the survey and they were on antibiotics. Among the 550 *E. coli* isolates from 550 children, 90% percent (n = 495) of the isolates were resistant to at least one of the commonly used antibiotics (ADR). However, multidrug resistance (MDR) was found in one-fourth of the isolates (41%, n = 226). Only 3% of isolates (n = 17) were found to be ESBL producers.

The resistance to single antibiotics and the combination of antibiotics (**Fig 3** and **Fig 4**) with demographic variables showed correlation with the resistance pattern (**Fig 5**). Commensal *E. coli* from males and females showed a similar pattern of resistance to all antibiotics except Ampicillin. Isolates with resistance to Ampicillin was higher in case of female children (57.14%) as compared to male (35.85%) but it was statistically non-significant. Isolates from children living in nuclear families were more resistant to ampicillin (OR 0.18, 95% CI: 0.11 – 0.28, p <0.01), ciprofloxacin (OR 0.48, 95% CI: 0.28 - 0.80, p <0.01), imipenem (OR 0.28, 95% CI: 0.10 - 0.80, p ≤ 0.01), and streptomycin (OR 0.31, 95% CI: 0.20 - 0.50, p <0.01), Whereas isolates from children living in joint families had less resistance to above antibiotics. *E. coli* from mothers having the education up to graduate level were less resistant to cefoxitin (OR 1.69, 95% CI: 1.16 – 2.46, p <0.01), ceftazidime (OR 0.75, 95% CI: 0.55 - 1.02, p < 0.02), imipenem (OR 3.24, 95% CI: 1.46 - 7.17, p < 0.01), and tetracycline (OR 1.75, 95% CI: 1.12 - 2.74, p <0.01). Paternal education was not associated with the resistant pattern of *E. coli* isolates from children.

**Fig 3:**
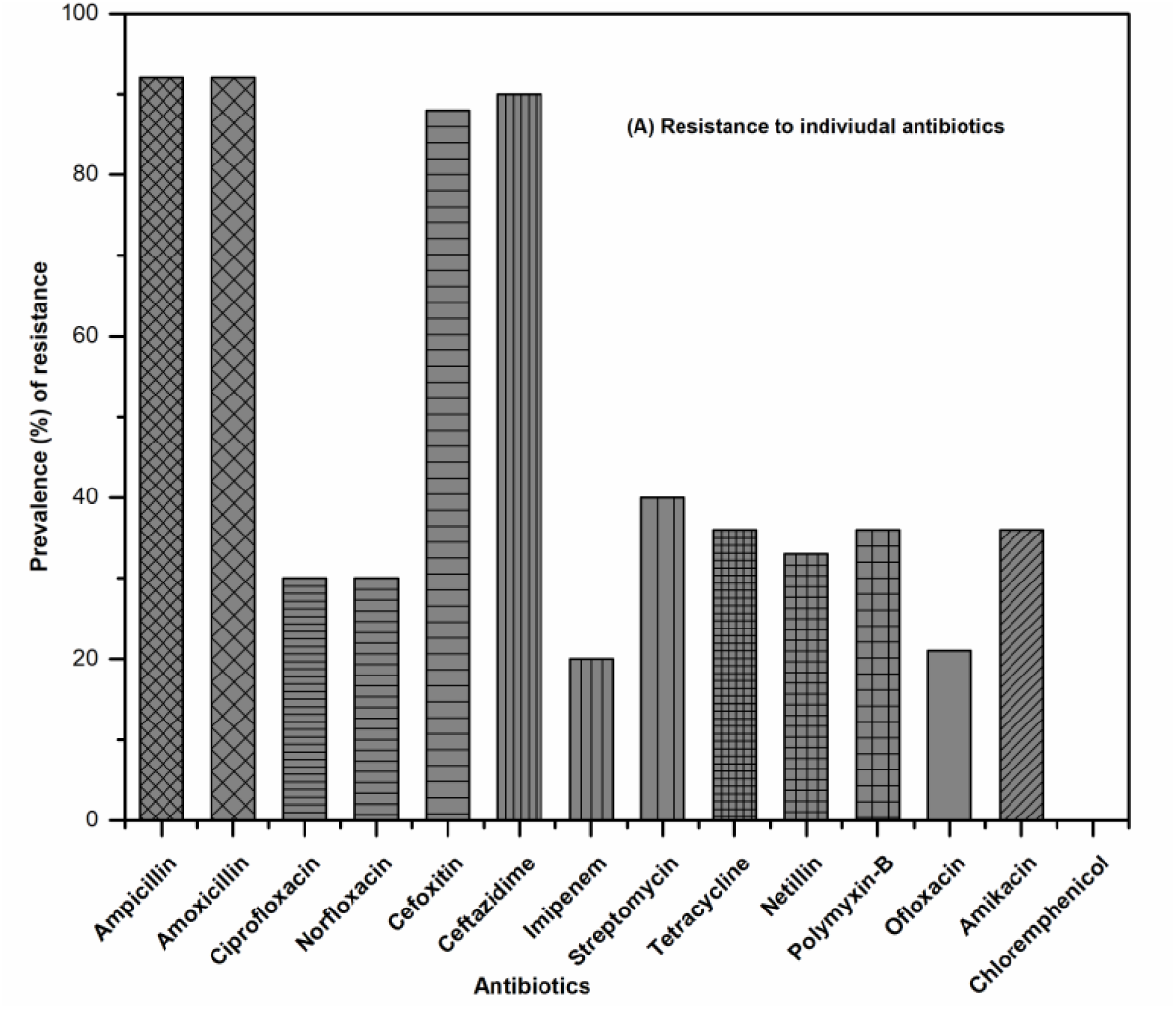
Resistance Pattern of *E. coli* isolates against individual antibiotics.

**Fig 4:**
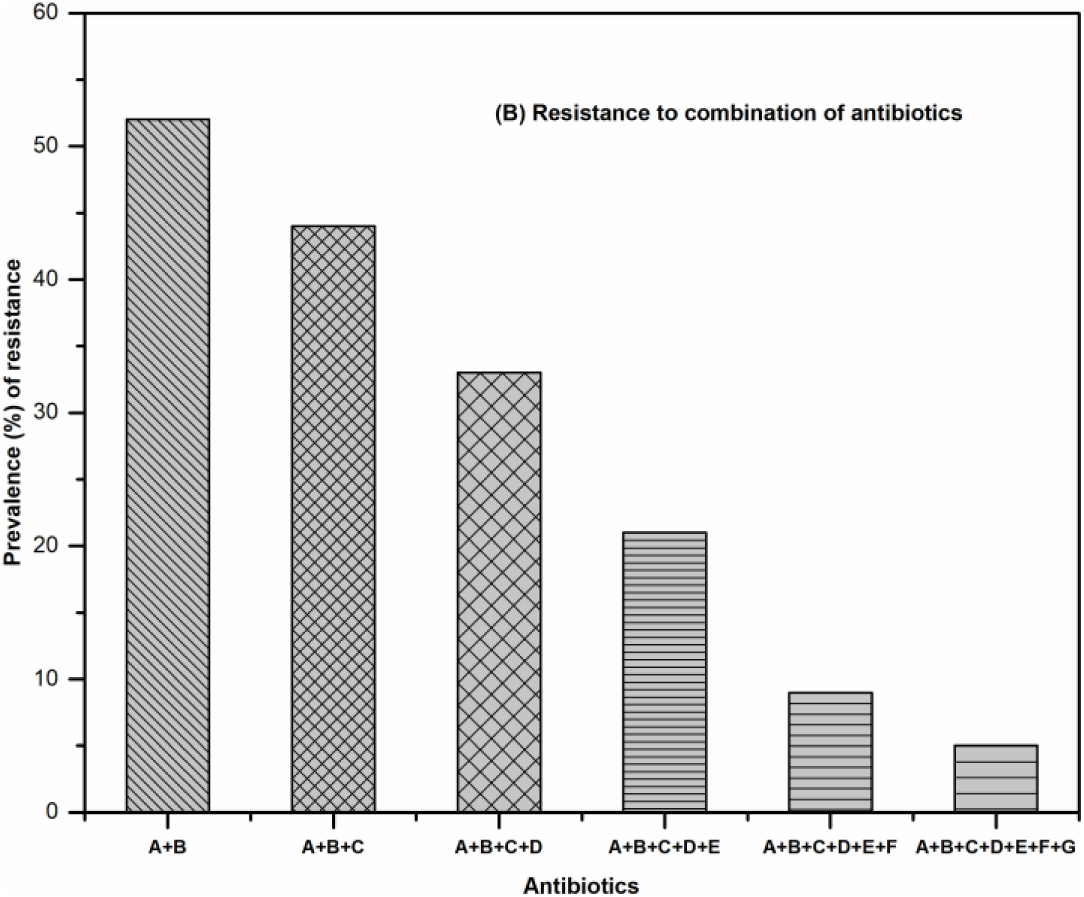
Resistance Pattern of *E. coli* against combination of different antibiotics. (Group A = Penicillin, B = Quinolones/Fluoroquinolones, C = Cephalosporin, D = Carbapenem, E = Aminoglycosides, F = Tetracycline, G = Polypeptide)

**Fig 5:**
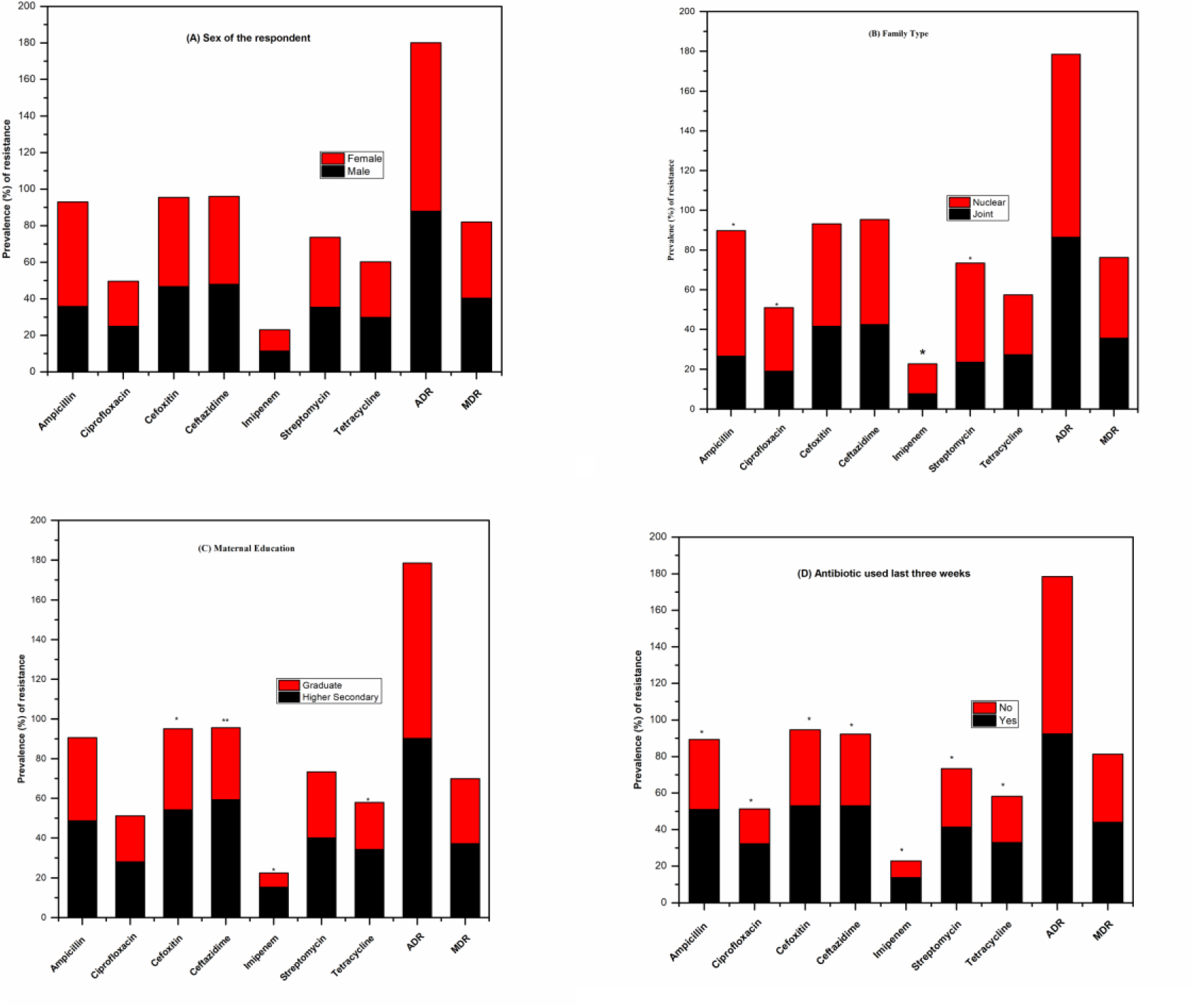
Prevalence (%) of resistance of *E. coli* to selected antibiotics associated with demographic variables. Prevalence (%) of resistance of *E. coli* to selected antibiotics associated with demographic variables. Resistance in *E. coli* isolated from (A) male and female (B) children living in joint and nuclear families (C) children with their mother having education up to graduate and higher secondary level and (D) children with prior antibiotic exposures. ^*^Statistically significant by chi-square analysis (p< 0.01); ^**^ statistically significant by multivariate logistic Regression.

ADR *E. coli* varied widely among villages from 30 to 90% and MDR *E. coli* varied from 10 to 41%. Children on antibiotics with a history of gastrointestinal illness (last three weeks) had a high rate of carriage of *E. coli* with ADR and MDR in comparison to children without any history of gastrointestinal illness and antibiotic use (**Fig 5**). However, these results of ADR and MDR were statistically not significant. Children receiving antibiotics in last three weeks had odds of carrying *E. coli* which were resistant to all the antibiotics and results are significant with p <0.01. Age had no association with ADR or MDR. Other demographic variables like economic status, caste, number of family members, paternal education, paternal occupation and maternal occupation were not associated with the resistant isolates.

In cluster analysis (Software: R Statistics, Package: ggplot, Function: heatmap.2), demographic data formed three distinct group based on the resistant pattern of *E. coli* isolates. The first group was formed by the *E. coli* isolates of male and female children. Though they showed high resistance pattern, it was statistically not significant (**Fig 5**) defining the AR in commensal *E. coli* is gender independent (**Group. 1; Fig. 6)**. Isolated *E. coli* from the children living in nuclear families having prior exposure to antibiotics and children of a mother who are school dropout showed highest resistance pattern (**Fig 5**) formed the second group (**Group 2; Fig 6**). The third group was formed by the children from the mothers who are graduates, without any prior antibiotic consumption before the study (last three weeks) and children living in a joint family **(Group 3; Fig 6)**. The third group was the carrier of least resistant *E. coli* (**Fig 6**). Therefore, analysis of cluster pattern showed a clear demographic association of AR in *E. coli* isolates. Antibiotic resistance pattern is independent of the gender of the children while they are highly dependent on the mother’s education and family type. Mother with higher education (graduate) and children living in the joint family are less like to carry resistant *E. coli* than those children living in nuclear families and mother having education till schooling level.

**Fig 6:**
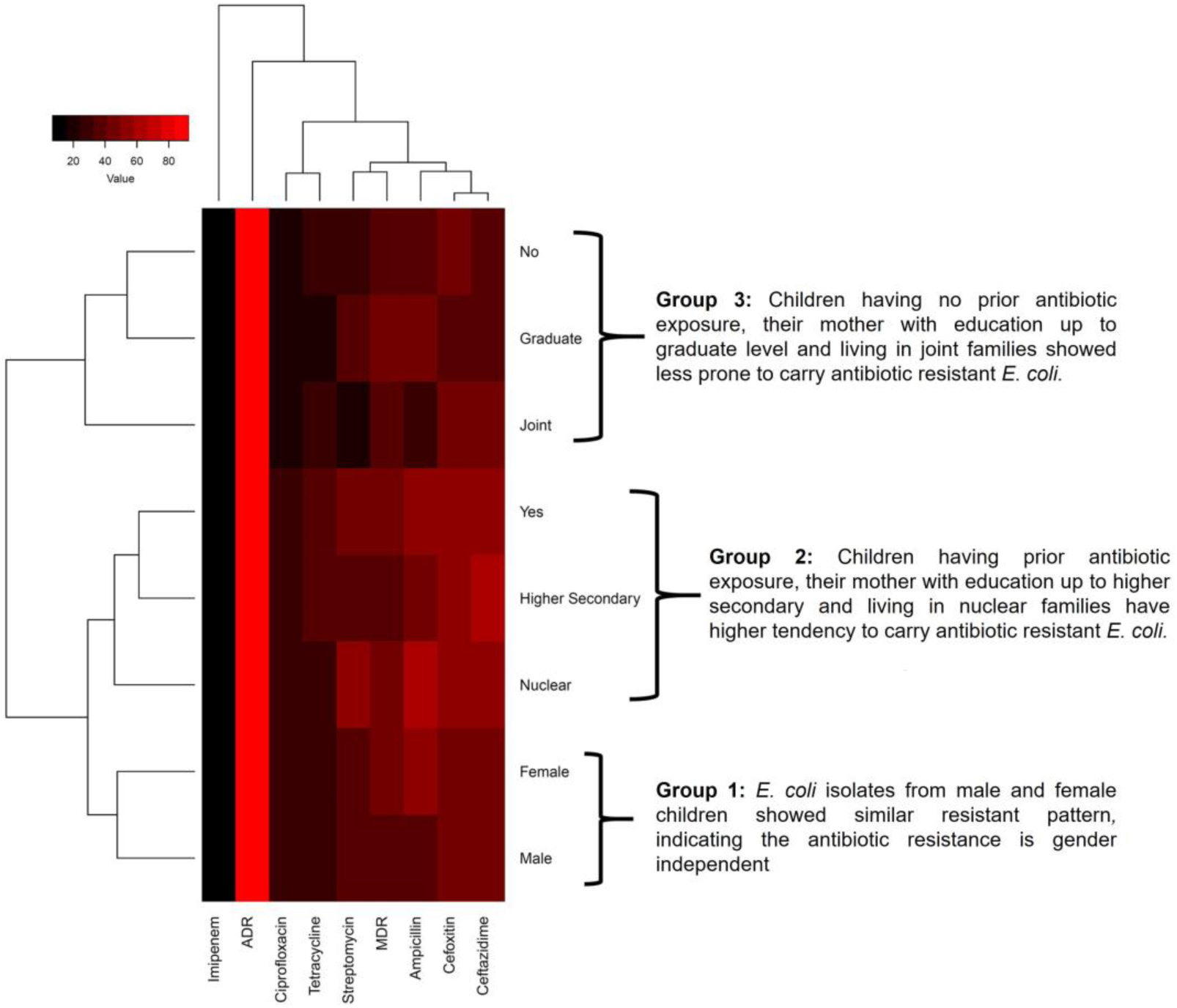
Cluster Analysis. Cluster analysis representing the relationship between demographic factors and pattern of antibiotic resistance. It formed three distinct groups/clusters based on the resistance pattern. Group 1 represents the *E. coli* isolates from male and female which showed similar resistance pattern. Group 2 showed the demographic factor which makes children more prone to carry higher resistant commensal *E. coli.* Group 3 showed the demographic factors for least resistance.

## Discussion

This is the first community-based study of its own to explore the prevalence of antibiotic-resistant commensal *Escherichia coli* in the children of rural areas of Sikkim as an indication or indicator of the emergence of antibiotic resistance. Antibiotic resistance (AR) has become one of the world’s most pressing public health problems of this era. Unnecessary use of antibiotics without prescription for treatment of common bacterial infection is considered as one of the leading causes of the emergence of antibiotic resistance in common pathogens in children and adults. The AR can create easily treatable illness to become dangerous infections leading to prolonged sufferings. It can spread to the family members, peer groups and community through different means. Shared distribution of *Escherichia coli* in the lower abdomen of human and susceptible nature to all the frequently used antibiotics makes it a good indicator bacterium to study the emergence of antibiotic resistance. The current study was designed to assess the prevalence and distribution of antibiotic resistance in *E. coli* isolated from the stool samples of the healthy children of Sikkim. The results were correspondingly correlated with demographic data to observe the effect of community structure on antibiotic resistance pattern of commensal *E. coli*.

### Antibiotic Resistant Patterns in Children in Sikkim

In this study, 90% of the isolated *E. coli* from the healthy children showed ADR property *i.e.* resistant to at least one antibiotic. This result is much higher than previously reported rates as observed in *E. coli* isolates from Central (72%), Eastern (38-68%) and Southern (63%) India [8][23][17]. A similar result was observed in the study conducted in Greece 1998 which reported that 40% of healthy children carried at least one antibiotic-resistant *E. coli* [23]. The study showed that bacteria carried by healthy children have commensal *E. coli* with higher resistance to the first generation of antibiotics (common antibiotics)like ampicillin (92%), streptomycin (40%), and tetracycline (36%). Correlative results were also reported in Peru and Bolivia where higher resistance was observed against common antibiotics (95%) [13]. One of the interesting results observed during the study was that as in Peru and Bolivia, high resistance against chloramphenicol (70%) was observed, while in Sikkim, chloramphenicol resistance was not found [13]. The probable reason behind the chloramphenicol susceptible isolates may be ascertained as less consumption of chloramphenicol because this antibiotic is used for diseases like typhoid, Cholera as well as Conjunctivitis which is less prevalent in Sikkim. The possible reason behind the development of resistance to the common antibiotics could be the frequent use or misuse of the antibiotics due to easy availability and affordability [8].

In Sikkim, 41% of the isolated *E. coli* showed MDR which is much higher than MDR reported from the eastern region of India. In the eastern region of India, Odisha had recorded 24% incidence of MDR *E. coli,* while in the middle part of India, Ujjain (Largest city of Madhya Pradesh) had recorded 33% of MDR *E. coli* isolates [8][23]. Southern India had recorded significantly higher percentage of commensal MDR *E. coli* (42%) than the eastern and middle India, but reciprocal to Sikkim (41%) [16]. The increased resistance pattern indicates the emergence of the resistant gene pool in commensal *E. coli* in India. This resistant pattern is a matter of concern as it is directly correlated with the health index of children of the community.

### Demographic factors affecting the resistance pattern

Demographic structure of a community is generally held as a key factor for emergence and outbreak of antibiotic resistance in bacteria. Children’s age, sex, parental education, socioeconomic status, family type and frequency of antibiotics taken are some of the important demographic parameters which were considered in the current study. During the study, we did not find any indication of the effect of age on the antibiotic resistance (age: 1-14) [17][23][8][5][13]. There are only a few available reports about the effect of children’s gender in the prevalence of antibiotic resistance. Some of the earlier studies reported that male children harbor more antibiotic resistant *E. coli* isolates in comparison to female child [23][13][5], whereas some other researchers found the opposites [10][8] Similar to the earlier findings we have also recorded the antibiotic resistance pattern among the isolates is independent of children’s gender. Statistically, the resistance pattern was found non-significant among the isolates from male and female children’s.

Maternal education as a demographic factor plays an important role in the development of antibiotic resistance in a community [24][8]. In this study, it was found that children with graduate mothers are less prone to carry resistant commensal *E. coli* than the mother having elementary schooling (**Fig 5**). Most of the antibiotics resistance was found for the antibiotics like cefoxitin (OR 1.69, 95% CI: 1.16 – 2.46, p <0.01), ceftazidime (OR 0.75, 95% CI: 0.55 - 1.02, p < 0.02), imipenem (OR 3.24, 95% CI: 1.46 - 7.17, p < 0.01) and tetracycline (OR 1.75, 95% CI: 1.12 - 2.74, p <0.01). Some of the earlier studies also reported the association between maternal education and increase of antibiotic resistance [8][25][26][27]. The reason could be the mothers with higher education are less likely to give antibiotics to their children without proper prescription as compared to a mother with lower education. Similarly, the children living in the joint family were found to be less prone to carrying the resistant isolates of *E. coli* as compared to the nuclear families. As for a reason, it could be due to the proper medical guidance of the elderly of the families in antibiotic consumption which is not possible in nuclear families where the parents are much busy with their chores and duties.

Geographically, the land of Sikkim host mountain ranges with varied altitude (350-8500m). In this study, the data showed that distance between market and villages and altitude of a village play an important role in the development of antibiotic resistance. The villages nearer to the market area or town area are prone to carrying more resistance *E. coli* compared to villages with greater distances. Similar patterns were also followed by the altitude of the village where people have easy excess to the pharmacy for antibiotics. A study carried out by Shakya *et al.,* (2013) showed that private pharmacies in Ujjain district were placed along with the main roads and those people living nearby with easy reach to these pharmacies to get antibiotics showed more antibiotic resistant *E. coli* isolates [8]. Besides above stated contributory factors, environmental factors were also supposed to play an important role in the variation of the incidence rates of antibiotic resistant *E. coli* isolates between villages. In the rural areas of Sikkim, water is supplied directly from spring (surface water) to community outlets without any treatment. There is no centralized water supply system. So, exposure to the environmental contaminants may create variation in the water supply from village to village. The human effluent and agricultural runoff along with resistant bacteria act as contaminants and serves for development of antibiotic resistance in commensal bacteria [28][3]. The current study only considered the antibiotic resistance pattern displayed by commensal *E. coli* in the rural communities of hills. This study only focuses on the resistant pattern among commensal *E. coli* in the rural communities. Though a large number of antibiotics were used in this study to detect the resistant pattern, we only used one isolate from each child. So, there exists a chance that some resistance isolates might have been missed out and the true resistance rate might be an underestimate of the problem.

## Conclusion

The current study identified a high prevalence of antibiotic-resistant commensal *E. coli* in children of rural areas of Sikkim, North Eastern India. The resistance pattern was studied for individual antibiotics and also for the combination of two to three classes of antibiotics. There was a significant correlation between the prevalence of antibiotic resistance commensal *E. coli* in the children and the demographic factors like mother’s education, family type, access and use of antibiotics. However, the prevalence of antibiotic resistance in *E. coli* isolates was found to be independent of child’s gender. A significantly high number of ADR and MDR isolates suggests an early need for the commitment to ensure the rational use of antibiotics as much as possible. This is the first report from hills of the eastern Himalayan region regarding resistance profile of gut *E. coli* from children. It warrants the future threat and suggests immediate counteractive measurements including educating the community about the proper uses of antibiotics and different hygiene protocols to tackle the future problem.

### Ethical Clearance

Approval for this study was granted from the Institutional Ethical Committee of Sikkim University, Gangtok, Sikkim.

## Acknowledgement

The authors wish to thank all the participating Children and their Parents of the rural community of Sikkim for their cooperation and pleasing behavior during the survey. The authors also extend their sincere gratitude for providing sufficient samples to the study. The authors also wish to thank the State Institute of Rural Development for their helping hand during sample collection and survey work. The authors also like to thank all the faculty member, non-teaching staff of the Department of Microbiology, Sikkim University for their continuous support and help throughout the study.

## Supporting File Information

S1: Consent Form in Two Languages (Nepali and English).

S2: Format of Questionnaire used during the study.

## Notes

**Conflict of Interest:** Authors have no conflict of Interest.

## References

1. Calva JJ, Cerón C, Calva JJ, Cero C, Diseases I, Nacional I, et al. Antimicrobial resistance in fecal flora?: longitudinal community-based surveillance of children from urban Mexico. These include?: Antimicrobial Resistance in Fecal Flora?: Longitudinal Community-Based Surveillance of Children from Urban Mexico. 1996;40: 1699–1702.

2. Laxminarayan R, Chaudhury RR. Antibiotic Resistance in India: Drivers and Opportunities for Action. PLoS Medicine. 2016;13:1–7. doi:10.1371/journal.pmed.1001974

3. Sahoo KC, Tamhankar AJ, Johansson E, Lundborg CS. Antibiotic use, resistance development and environmental factors: a qualitative study among healthcare professionals in Orissa, India. BMC Public Health; 2010. 2010; 10: 1–10. doi:10.1186/1471-2458-10-629

4. Wright GD. The antibiotic resistome: the nexus of chemical and genetic diversity. Nature Reviews Microbiology. 2007;5: 175–186. doi:10.1038/nrmicro1614

5. Bartoloni A, Cutts F, Leoni S, Austin CC, Mantella A, Guglielmetti P, et al. Patterns of antimicrobial use and antimicrobial resistance among healthy children in Bolivia. Trop Med Int Health. 1998;3: 116–123. Available: http://onlinelibrary.wiley.com/store/10.1046/j.13653156.1998.00201.x/asset/j.13653156.1998.00201.x.pdf?v=1&t=ii8u6tke&s=861bdf4354f1526f24cb97a6e96c56dc2b806c352b806c35

6. Takahashi K. Interaction between the Intestinal Immune System and Commensal Bacteria and Its Effect on the Regulation of Allergic Reactions. Bioscience, Biotechnology, and Biochemistry. 2010;74: 691–695. doi:10.1271/bbb.90962

7. Silva N, Igrejas G, Gonçalves A, Poeta P. Commensal gut bacteria: Distribution of Enterococcus species and prevalence of Escherichia coli phylogenetic groups in animals and humans in Portugal. Annals of Microbiology. 2012;62: 449–459. doi:10.1007/s13213-011-0308-4

8. Shakya P, Barrett P, Diwan V, Marothi Y, Shah H, Chhari N, et al. Antibiotic resistance among Escherichia coli isolates from stool samples of children aged 3 to 14 years from Ujjain, India. BMC Infectious Diseases. 2013;13: 1–6. doi:10.1186/1471-2334-13-477

9. Lestari ES, Severin JA, Filius PMG, Kuntaman K, Duerink DO, Hadi U, et al. Antimicrobial resistance among commensal isolates of Escherichia coli and Staphylococcus aureus in the Indonesian population inside and outside hospitals. European Journal of Clinical Microbiology and Infectious Diseases. 2008;27: 45– 51. doi:10.1007/s10096-007-0396-z

10. Vatopoulos AC, Varvaresou E, Petridou E, Moustaki M, Kyriakopoulos M, Kapogiannis D, et al. High rates of antibiotic resistance among normal fecal flora Escherichia coli isolates in children from Greece. Clinical microbiology and infection?: the official publication of the European Society of Clinical Microbiology and Infectious Diseases. 1998;4: 563–569. doi:10.1111/j.1469-0691.1998.tb00038.x

11. Oluyege AO, Ojo-Bola O, Oludada OE. Carriage of antibiotic resistant commensal E. coli in infants below 5 months in Ado-Ekiti. International Journal of Current Microbiology and Applied Sciences. 2015;4: 1096–1102.

12. Infante B, Grape M, Larsson M, Kristiansson C, Pallecchi L, Rossolini GM, et al. Acquired sulphonamide resistance genes in faecal Escherichia coli from healthy children in Bolivia and Peru. International Journal of Antimicrobial Agents. 2005;25: 308–312. doi:10.1016/j.ijantimicag.2004.12.004

13. Bartoloni A, Pallecchi L, Benedetti M, Fernandez C, Vallejos Y, Guzman E, et al. Multidrug-resistant commensal Escherichia coli in children, Peru and Bolivia. Emerging Infectious Diseases. 2006;12: 907–913. doi:10.3201/eid1206.051258

14. World Health Organization, Ministry of Health & Family Welfare G of I. National Action Plan on Antimicrobial Resistance (NAP-AMR) 2017-2021 [Internet]. 2017. Available: http://www.nicd.nic.in/writereaddata/mainlinkfile/File645.pdf

15. Van Boeckel TP, Gandra S, Ashok A, Caudron Q, Grenfell BT, Levin SA, et al. Global antibiotic consumption 2000 to 2010: An analysis of national pharmaceutical sales data. The Lancet Infectious Diseases. Elsevier Ltd; 2014;14: 742–750. doi:10.1016/S1473-3099(14)70780-7

16. Mathai E, Chandy S, Thomas K, Antoniswamy B, Joseph I, Mathai M, et al. Antimicrobial resistance surveillance among commensal Escherichia coli in rural and urban areas in Southern India. Tropical medicine & international health?: TM & IH. 2008;13: 41–45. doi:10.1111/j.1365-3156.2007.01969.x

17. Seidman JC, Anitha KP, Kanungo R, Bourgeois AL, Coles CL. Risk Factor for antibiotic-resistant E coli in children in a rural aeas. epidemiology infection. 2009;137: 879–888. doi:10.1002/ana.22528.Toll-like

18. Tambe S, Kharel G, Subba S, Arrawatia ML. Rural Water Security in the Sikkim Himalaya?: Status, Initiatives and Future Strategy. 2012.

19. Issues R, Management W. Source?: Census of India 2011 Data State?: Sikkim Sikkim Population 2011 Sikkim Literacy Rate 2011 Sikkim Density 2011. 2011.

20. CensusofIndia:Sikkim. District Census Handbook: North, West, South And East Districts Village And Town Wise Primary Census Abstract (PCA). Dirctorate of Census Operations: Sikkim. 2011; 1–308. Available: http://www.censusindia.gov.in/2011census/dchb/1100_PART_B_DCHB_SIKKIM.pdf

21. Sullivan KM, Dean A. OpenEpi: A Web-based Epidemiologic and Statistical Calculator for Public Health. Public Health Reports. 2009;124: 471–474. doi:10.2307/25682255

22. Clinical and Laboratory Standards Institute. Performance standards for antimicrobial disk susceptibility tests: approved standard [Internet]. 10th ed. CLSI. Wayne, PA, USA; 2009. doi:M02-A11

23. Sahoo KC, Tamhankar AJ, Sahoo S. Geographical Variation in Antibiotic-Resistant Escherichia coli Isolates from Stool, Cow-Dung and Drinking Water. International journal of environmental research and public health. 2012;9: 746– 759. doi:10.3390/ijerph9030746

24. Allin S, Stabile M. Socioeconomic status and child health: what is the role of health care, health conditions, injuries and maternal health? Health Economics, Policy and Law. 2012;7: 227–242. doi:10.1017/S174413311100034X

25. Bi P, Tong S, Parton KA. Family self-medication and antibiotics abuse for children and juveniles in a Chinese city. Social Science & Medicine. Pergamon; 2000;50: 1445–1450. doi:10.1016/S0277-9536(99)00304-4

26. Okumura J, Wakai S, Umenai T. Drug utilisation and self-medication in rural communities in Vietnam. Social Science & Medicine. Pergamon; 2002;54: 1875– 1886. doi:10.1016/S0277-9536(01)00155-1

27. Togoobaatar G, Ikeda N, Ali M, Sonomjamts M, Dashdemberel S, Mori R, et al. Survey of non-prescribed use of antibiotics for children in an urban community in Mongolia. Bulletin of the World Health Organization. World Health Organization; 2010;88: 930–936. doi:10.1590/S0042-96862010001200014

28. Sabde YD, Diwan V, Saraf VS, Mahadik VK, Diwan VK, De Costa A. Mapping private pharmacies and their characteristics in Ujjain district, Central India. BMC Health Services Research. BioMed Central Ltd; 2011;11: 351. doi:10.1186/1472- 6963-11-351

